# Season-dependent processing of innate conspecific vocalizations in the male and female European starling (*Sturnus vulgaris*)

**DOI:** 10.1101/2023.07.20.549888

**Authors:** N.D. Vidas-Guscic, E. Jonckers, J. Van Audekerke, J. Orije, J. Hamaide, G. Majumdar, M. Verhoye, A. Van der Linden

**Affiliations:** Bio-Imaging Lab, University of Antwerp, Antwerp Belgium

**Keywords:** European starling, begging calls, caudomedial nidopallium, auditory perception, Field L, functional magnetic resonance imaging, songbird

## Abstract

Avian innate nestling begging calls are similar to human infant cries in the behavioral response they elicit. However, it remains unknown whether the auditory processing of innate begging calls changes in seasonal songbirds from non-breeding to breeding season when hormonal neuromodulation of the auditory forebrain occurs.

An fMRI experiment was set up to expose male and female European starlings (*Sturnus vulgaris*) to recordings of seasonal conspecific nestling begging calls in the breeding and non-breeding season. This response was compared with their response to conspecific warble motifs and artificial pure tones, both proven seasonally invariable at least in the male starling’s neural response.

Our results demonstrate significant seasonal variation in auditory forebrain responses exclusively elicited by begging calls and not by the applied control stimuli. Right Field L and the Caudomedial Nidopallium (NCM) seemed, irrespective of season or sex, more sensitive in response to begging than to control stimuli. A seasonal differential response specifically to begging calls was found in both sexes in a ventral midsagittal region of NCM.

Our findings thereby support the functional fine-tuning of vocal communications between sender and receiver in a breeding context for innate vocalizations and are in line with the bi-parenting behavior in this species.

## Introduction

Call-type innate vocalizations in songbirds have received far less attention from scientists investigating the neural substrate of vocal communication. This is mainly because songbirds’ learned songs develop very similarly to human speech, making them an excellent model for vocal learning (Marler, 2004; Bolhuis, Okanoya and Scharff, 2010). Over the past decades, several studies in humans have attempted to visualize the neural substrate for the processing of innate human infant cries relating these findings to parental care (Lorberbaum *et al*., 1998; Swain, 2008; Piallini, Palo and Simonelli, 2015; Kim *et al*., 2017). Avian innate nestling calls are comparable to human infant cries in the behavioral response they elicit. However, to our knowledge, no study has investigated whether the auditory processing of the nestling calls is different during the breeding as compared to the non-breeding season in birds. However such investigations can shed light on the neuromodulatory impact of reproductive hormones on the auditory sensitivity to offspring’s vocalization and its link with proper parental care.

Seasonal songbirds such as the European starling (*Sturnus vulgaris*) display a photoperiod-sensitive development of the gonadal system resulting in a tight synchronization between hatching and hatchling’s food supply (for review see: Dawson et al., 2001; Gwinner, 2003). Most breeding-related behaviors such as singing and nest building are also energetically costly and therefore restricted to the breeding season (Daan *et al*., 1989). This seasonal breeding requires dedicated strategies to cope with the changing behavioral relevance of communication signals and neuromodulation of the auditory system by steroid hormones could play an important role in this seasonal finetuning. Starling’s learned songs, classified into individual- and species-specific songs, are both used in- and outside the breeding season by both sexes, but their abundance and relevance change between the seasons (Hausberger, 1997) (George *et al*., 2008). Using auditory functional MRI (fMRI) in male starlings, we could confirm the seasonal relevance of specific types of vocalizations as reflected in the neural activity of a secondary auditory brain region: the caudomedial nidopallium (NCM) (De Groof et al., 2013) and even demonstrate that the seasonal shift in auditory attention was mediated by local changes in estrogen (De Groof et al, 2017).

Like pair formation in the breeding season, also the introduction of nestlings changes the social group structure. As an altricial species, European starling nestlings possess a repertoire of innate vocalizations at birth through which they solicit parental attention and care, such as *alarm calls* for defense and the nestling-derived food *begging calls* for nourishment. Begging calls are directed at both parents and both parents take part in feeding the nestlings in starlings (Corney and Barber, 2018). Their bi-parental feeding strategy indicates that begging calls should be perceived with equal importance by both parents. Also, because it is a seasonally reproducing songbird, exposure to nestling begging calls is only expected to occur during the breeding season. This is different from the different types of starling songs that are communicated throughout the seasons. The question remains whether recordings of this innate vocalization in the absence of hatchlings will be perceived differentially between breeding and non-breeding season and thus whether the bird can functionally classify these vocal signals based on their seasonal relevance. In songbirds, the NCM and caudomedial mesopallium (CMM) are auditory regions analogous to the human secondary auditory cortex and Field L is homologous to the human primary auditory cortex (Mello, Vicario and Clayton, 1992; Nagel *et al*., 2011). Seasonal dynamics have been reported in the NCM for the perception of *learned* vocalizations both in song sparrows (Caras et al., 2012) and starlings (De Groof et al., 2013, 2017).

The present work will investigate neural activations in adult male and female starlings that have not built a nest using auditory fMRI upon recorded nestling begging call stimulation as compared to control sounds (starling warble song fragments and pure tones) under anesthesia. Auditory fMRI is a non-invasive imaging technique that makes it possible to visualize differential neuronal responses to different stimuli presented repeatedly over time in the same subject. In this study, we aim to answer the following questions: 1) which regions of the avian auditory forebrain are responsible for perceiving begging calls, and is this activation higher or lower as compared to control stimuli, 2) does this activation pattern change from breeding to non-breeding season and 3) is this region and the intensity of the auditory neuronal activation the same for both sexes. The outcome will provide more insight into how the brain deals with auditory signals while their social relevance is changing and whether this is different for vocalizations that are learned or innate.

## Material and methods

### Ethics statement

All procedures and animal handling were performed in accordance with the European guidelines for the care and use of laboratory animals (2010/63/EEC) and approved by the Committee on Animal Care and Use at the University of Antwerp, Belgium (ECD # 2018–88).

### Subjects

Eleven adult male and nine adult female European starlings (85 ±10 grams) (*Sturnus vulgaris*) were used in this experiment. Birds were wild caught in Cyprus in January 2018/2019. The population was divided into two sex-mixed groups and housed in two indoor aviaries (1.40 × 2.20 × 2.10 m) of the Bio-Imaging Lab at the University of Antwerp (Antwerp, Belgium). No nest boxes were provided, hence no nesting behavior was observed. Food and water were available *ad libitum* in the aviaries.

### Photoperiodic manipulations to induce seasonality

To investigate whether begging calls are categorized differently based on seasonal behavioral relevance, we controlled seasonality by an artificial light-dark cycle, an approved method to induce photostimulation (breeding) and photorephractoriness (non-breeding) (Dawson *et al*., 2001). This regime consisted of 10 weeks of short days (SD: 8 hours light) followed by 16 weeks of long days (LD: 14 hours light), which successfully simulated natural photoperiodicity at an accelerated rate (Bernard and Ball, 1997; De Groof et al., 2013, 2017) (**figure 1**).

**Figure 1.**
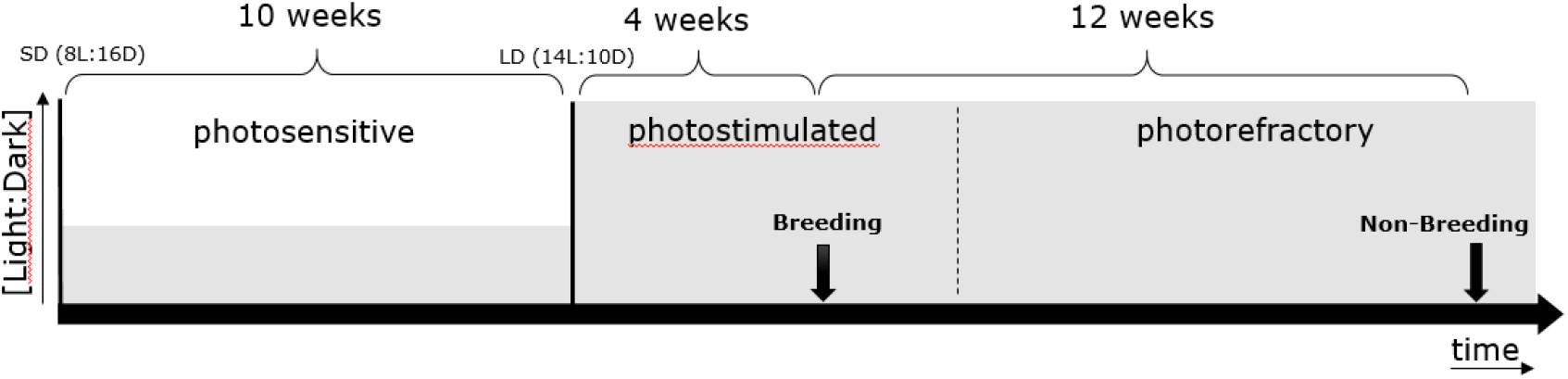
Schematic representation of photoperiodic manipulation to induce photostimulation and photorefractoryness. Arrows indicate fMRI scanning periods (one in breeding and one in non-breeding periods). Solid lines mark clear transitions between photostages, whereas the dotted line represents a gradual transition. Grey levels indicate the [light:dark] ratio.

Birds were made photosensitive in anticipation of the experiments by shifting them to SD (**figure 1**). After 10 weeks of SD conditions, birds are photosensitive and ready to respond to LD stimulation. The subsequent shift to LD induces the seasonal development of their gonads and song control system plasticity to mimic the breeding season (=**photostimulated**). After approximately four weeks of LD exposure, starlings reached peak photostimulation. After another 12 weeks of LD starlings lost their sensitivity to long daylight periods (=**photorefractory**) as indicated by post-nuptial molting, which occurs when gonads are regressing and plasma testosterone levels drop (Gwinner, 1977) (Dawson, 2003).

The starling’s brain activity was visualized using auditory fMRI after four weeks in the artificial breeding and after twelve weeks in the non-breeding periods (see **figure 1**).

### Anesthesia and physiology

In preparation for the MRI imaging session, birds were individually retrieved from their home cage in a transportation box to reduce external auditory and visual stimulation and minimize confounding factors that could influence neural responsivity during the fMRI scans. Additionally, they were then kept in sensory-reduced conditions for at least thirty minutes before scanning.

Animals were anesthetized with a 0.2 ml intramuscular bolus injection in the pectoral muscle containing a mixture of medetomidine (10 ml, 1 mg/ml Domitor, Pfizer, Germany) and ketamine (0.5 ml, 50 mg/ml Anesketine, Eurovet Animal Health, the Netherlands). Anesthesia was maintained through continuous intramuscular infusion with the same anesthetic mixture at a rate of 0.12 ml/h. Minutes after injection, consciousness was assessed with the toe pinch test before the birds were restrained in the MRI scanner in a prone position.

During scanning, animals were breathing a mixture of oxygen and nitrogen (200 and 400cm^3^/min) delivered through the beak-holder. The breathing rate was followed with a pneumatic sensory pad underneath the animal’s chest (SA Instruments Inc., US).

Body temperature was maintained at 40 ±0.5°C using a cloacal temperature probe, connected to a feedback-controlled air heating system (SA Instruments Inc., US). After scanning, birds were administered with an intramuscular bolus of 0.2 ml Atipamezole (0.5mg/ml Antisedan, Zoetis, US) to reverse the effects of medetomidine.

### Auditory stimulation

Magnetless dynamic speakers (Visaton, Germany) connected to a desktop featuring Presentation V18.3 (Neurobehavioral Systems Inc., US) in the control room were used as stimulation device. Between different subjects’ scans, the left and right headphones were switched to account for any hemispheric bias in auditory stimulation resulting from potential speaker inequality. Song (individual warbles type 3A) and begging fragments were obtained from the Animal and Human Ethology research group of the University of Rennes (Rennes, France) and were thus unfamiliar to the starlings in this experiment. Type 3A individual warbles were selected as naturalistic control sound because previous work has demonstrated that the response to this type of stimulation did not change between seasons as it contains information that is important all year round (De Groof et al., 2013). Artificial *pure tones* (stimulus made of 3 and 7 kHz, interleaved with 0.5s of silence) were also included as a control to exclude potential seasonal changes in auditory acuity (**figure 2 A-C**). The intensities of the different stimuli were normalized to 67 decibels with Praat V6.0.50 software (Paul Boersma, University of Amsterdam).

**Figure 2.**
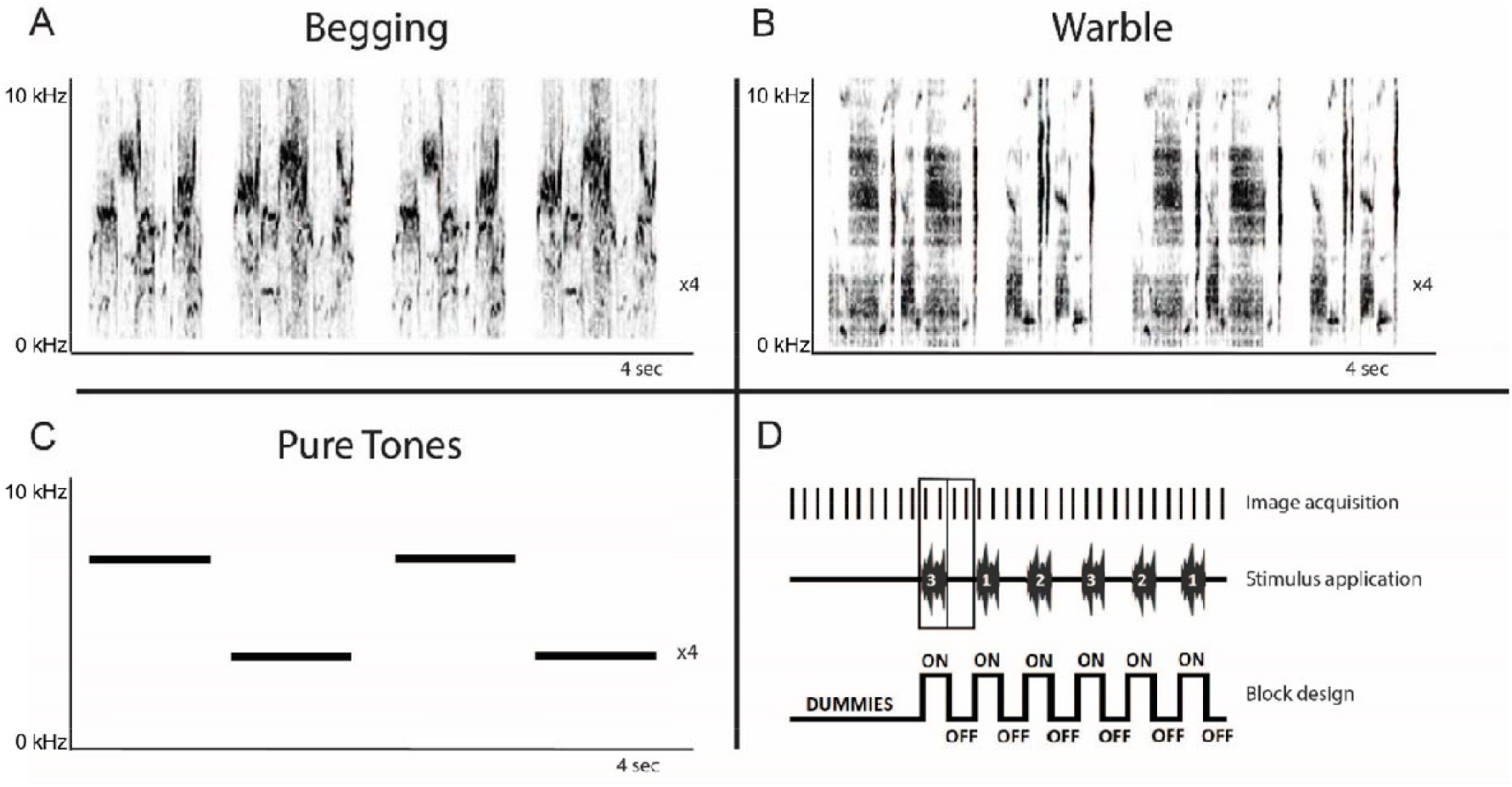
Overview of sonograms and acquisition paradigm. **A)** Begging call: monosyllabic vocalization that is loud and relatively long for a call and emitted in bouts directed at parents that are present at the nest (Elie and Theunissen, 2015) **B)** Individual warbles Class 3A: initial motifs taken from a quiet warbling song (De Groof et al., 2013). **C)** Pure tones: 3 and 7kHz artificial control sounds. **D)** Schematic representation of the initial part of a randomized block design with each ON-block representing the acquisition of two BOLD fMRI images during auditory stimulation, interweaved by OFF-blocks consisting of two fMRI images acquired during rest (baseline).

The stimulus protocol consists of a randomized block design of 16s ON / 16s OFF blocks. Each stimulation ON-block consisted of two similar sound fragments that were presented in an ABAB pattern with 0.5s rest between every fragment. This pattern was repeated four times, which resulted in a total stimulus duration of 16s per ON-block, followed by a complementary OFF-block of 16s. Each stimulation block was presented 21 times per stimulus (Begging call, individual warbles 3A, Pure Tones), and two fMRI images were recorded per stimulation (ON or OFF) block, resulting in 252 MRI images. Ten dummy repetitions without stimulation were included at the start of the fMRI scan to allow for magnetization stabilization, giving a total of 262 MRI images per scan (**figure 2D**).

### Image acquisition

All MRI measurements were performed on a PharmaScan 70/16 USR horizontal MR system (Bruker, Germany) equipped with a volume transmit coil and a four-channel parallel receive array coil (Bruker, Germany). To investigate changes in the Blood Oxygen Level Dependent (BOLD) signal over time, T2-weighted turbo-RARE (Rapid Acquisition Relaxation Enhanced) images were acquired with the following parameters: Effective TE= 50.60ms, TR= 2000ms, RARE factor= 8. Fifteen sagittal whole brain slices were recorded with a ventral-dorsal read orientation, a slice thickness of 1.0mm, and a 0.08mm slice gap. The matrix size was [64×32] with a field of view of (27×27)mm^2^ giving an in-plane resolution of (0.33×0.67)mm^2^. Additionally, fat suppression was enabled, and saturation slices were used to remove signals originating from adipose tissue in the neck and for rapid eye movements. Functional image time series were reconstructed with a trapezoid filter (0.25 × 0.75) in the frequency and phase encoding direction.

An fMRI scan was considered successful if (*1*) a significant BOLD response could be detected in the auditory forebrain at the exploratory threshold P_uncorrected_<0.05, (*2*) framewise displacement was no more than two voxels and (*3*) no artifacts were present in the images. Unsuccessful scans were repeated after a minimum of two days, allowing birds to recuperate from anesthesia.

### Image processing

The raw datasets were acquired with ParaVision 6.0.1 (Bruker, Germany) and converted to the nifti file format, using an in-house script in MATLAB R2017b (MathWorks, US). Statistical Parametric Mapping 12 (SPM12) (FIL methods group, University College London) was used to align all functional images to the first image, based on a six-parameter (rigid body) spatial transformation. A mixed-sex population-based template was created from the first repetitions of each fMRI scan in Advanced Normalization Tools (ANTs). Next, fMRI scans of all subjects and all seasons were registered to this study-based template in SPM using a 12-parameter affine transformation, followed by non-linear deformations. Finally, the data were smoothed in-plane using a Gaussian kernel with an FWHM of (0.66 × 1.34) mm^2^. We applied a high pass filter (352 seconds) to remove low-frequency drift in the BOLD signal. Next, for each subject, the BOLD signal in each voxel was modeled with a Finite Impulse Response function. A starling MRI atlas (De Groof *et al*., 2016) was normalized to the study-based template, which functioned as a high-resolution anatomical reference. This atlas treats all nuclei as a single entity and cannot separate between functional subdivisions of the NCM and Field L. Hence, all conclusions will be based on approximations of delineations provided from the literature (Fortune and Margoliash, 1992; Pinaud *et al*., 2006).

### Statistical analysis

Statistical voxel-based analyses were performed for each scan using a mass-univariate approach based on the General Linear Model (GLM) implemented in SPM12. In each voxel, the significant BOLD response for the three stimuli was computed by digitally subtracting the OFF signal from the ON signal. *t*-contrasts (stimulus > rest) were defined for each stimulus separately and together, resulting in contrast files to be used in the next processing steps.

First, a one-sample t-test was performed on the first-level contrasts of all stimulations (begging call, individual warble, and pure tones) over rest blocks. The voxels that demonstrated significant activity over rest were used as a region of interest for small-volume correction in further analysis. Next, separate ANOVA models were designed to model the separate effects of sex, season, and stimulus, and finally, a grand design containing all factors to model potential interaction effects in a three-way ANOVA was used.

The familywise error (FWE) correcting method using random field theory that is built-in SPM12 was used to correct for false positives. Only findings with a cluster-wise error rate of *P*_*FWE*_ < 0.05 that was larger than ten voxels (k_*e*_ >10) were considered statistically significant.

## Results

A longitudinal auditory fMRI study was set up to visualize and quantify the neural responses elicited in male and female starlings upon exposure to juvenile begging calls, which are innate by nature and have a strong differential seasonal relevance. Hereto, seasonal songbirds were brought subsequently into artificial breeding and non-breeding conditions to assess the effects of seasonality and sex on their auditory processing.

### Begging Calls produce stronger bilateral activation in the auditory forebrain than control sounds regardless of seasonality or sex

Firstly, one sample t-tests were performed on a group level for each stimulus over baseline to confirm the responsiveness to all stimuli and to assess the topography and the relative response amplitude of the BOLD response to each stimulus, regardless of seasonality or sex. This revealed activations to all stimuli in Field L, NCM, and CMM (**Supplementary figure 1**; P_UNC_<0.001, T_max_=9.24 and k_e_=69 voxels). This activation upon all stimuli over rest was used as a mask for small volume correction in the subsequent voxel-based statistics. Subsequently, one sample t-test for each separate stimulus compared to the rest block demonstrated activation in the bilateral auditory nuclei Field L (primary auditory region), NCM, and CMM (secondary auditory regions) for every stimulus (Figure 3A-C).

**Figure 3.**
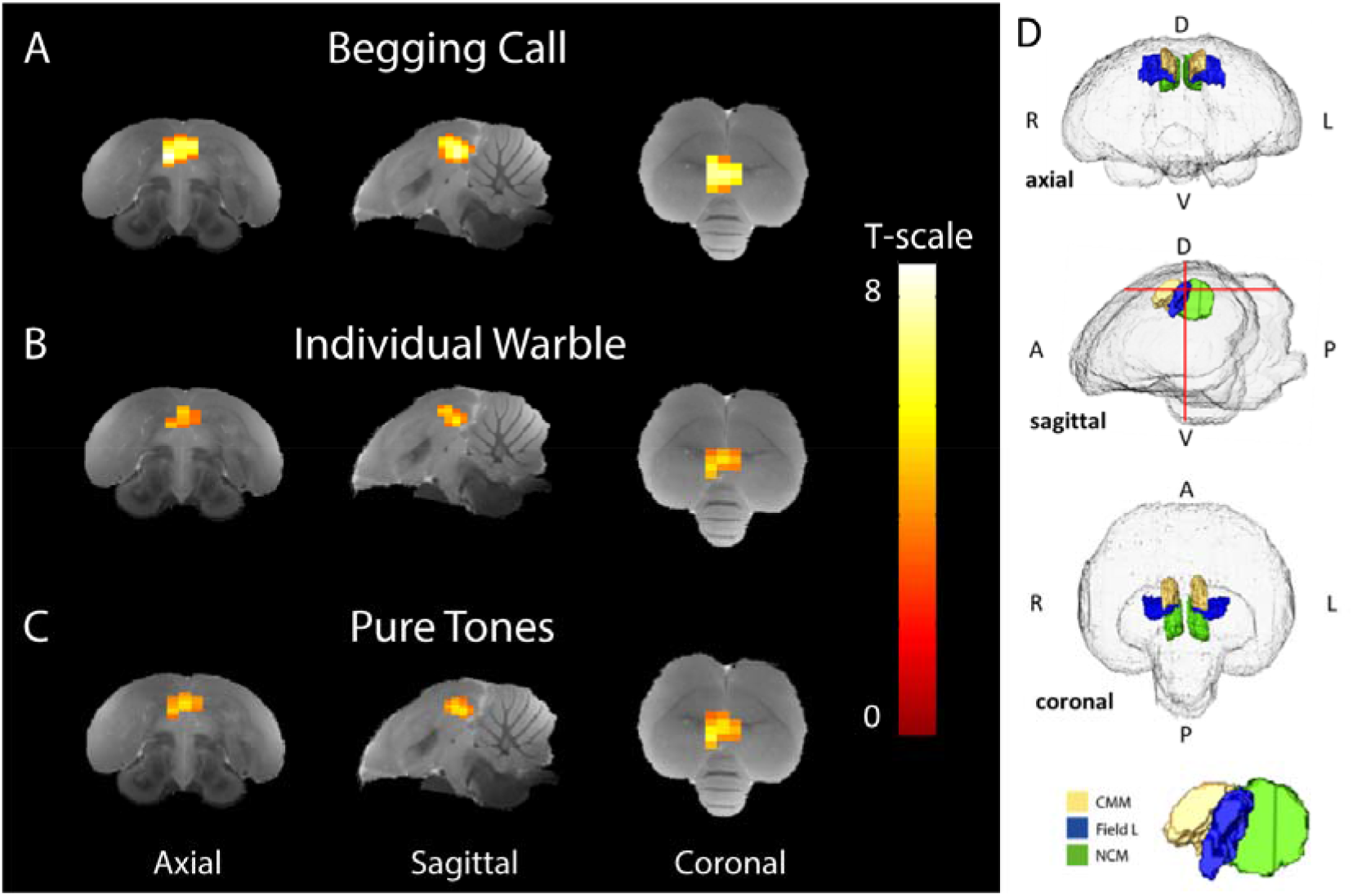
Average stimulus-specific neural activation (breeding and non-breeding season in both sexes) higher than rest periods. Voxels showing BOLD signal responses that are significantly stronger during stimulation than rest blocks have been superimposed on the starling MRI atlas (De Groof et al., 2016) for **(A)** begging calls, (**B)** individual warbles class 3A, and **(C)** Pure Tones (3 + 7 kHz). The color bar indicates the t-statistic for each voxel. **(D)** Transparent volume rendering of the avian brain containing color-coded volume renders of auditory forebrain structures with crosshairs indicating the slice orientation displayed in A-C and on the bottom, a 3D render of with yellow, blue, and green indicating CMM, Field L, and NCM respectively (One sample t-tests: P_Unc_<0.001, k_voxels_>10). D=dorsal, V=ventral A=anterior P=posterior, L=left, R=right.

Begging calls elicited more widespread BOLD activation compared to the warbles and pure tones, resulting in a larger cluster of 81 voxels in the auditory forebrain (P_FWE-peak_<0.0001, T_peak_=8.51) **figure 3A**). The individual warbles motifs produced smaller spatial activation than the begging call, resulting in a cluster of 45 voxels located in Field L, NCM, and CMM (P_FWE-peak_=0.002, T_peak_=5.83) (**figure 3B**). Pure tones of 3 and 7 kHz, elicited activation that was spatially and statistically similar to the individual warbles with a significantly higher BOLD response compared to baseline in a total of 53 voxels in Field L, NCM, and CMM (P_FWE-peak_<0.0001, T_peak_=6.33) **(figure 3C**). Overall, the begging calls activation is stronger and recruits a larger area in the auditory regions, with higher *t*-statistics as compared to the naturalistic and artificial control sounds.

### Begging Calls produce a differential activation in comparison to control sounds in right Field L and NCM with the highest difference in activation in right Field L

To pinpoint which voxels have a differential activation upon begging calls, individual warble, and pure tones, an ANOVA was performed to test the main effect of stimulus (**Figure 4A**). This revealed a cluster of 16 voxels in right Field L and NCM (P_FWE-peak_<0.0001, F_peak_=19.89). *Post-hoc* t-tests were performed to test if the amplitude of the bold response was significantly higher for the begging calls compared to the other stimuli. Therefore we statistically compared the relative signal increase over the baseline of the begging calls to the relative activation over the baseline for individual warbles (**Figure 4C)** and pure tones (**figure 4E**). For the begging calls compared to individual warbles, we observed significantly higher activation for begging calls in field L and NCM with the biggest difference in activation (white voxels in **figure 4**) in Field L in a cluster of 19 voxels (P_FWE-peak_<0.0001, T_peak_=5.72). For the comparison with pure tones, a cluster of 14 voxels was also found in Field L (P_FWE-peak_<0.0001, T_peak_=5.41). No other pairwise comparison of stimuli revealed voxels that produce significant activation, indicating that specifically, the begging calls produce stronger activation than the other stimuli in Field L and NCM with the highest significance in Field L.

**Figure 4.**
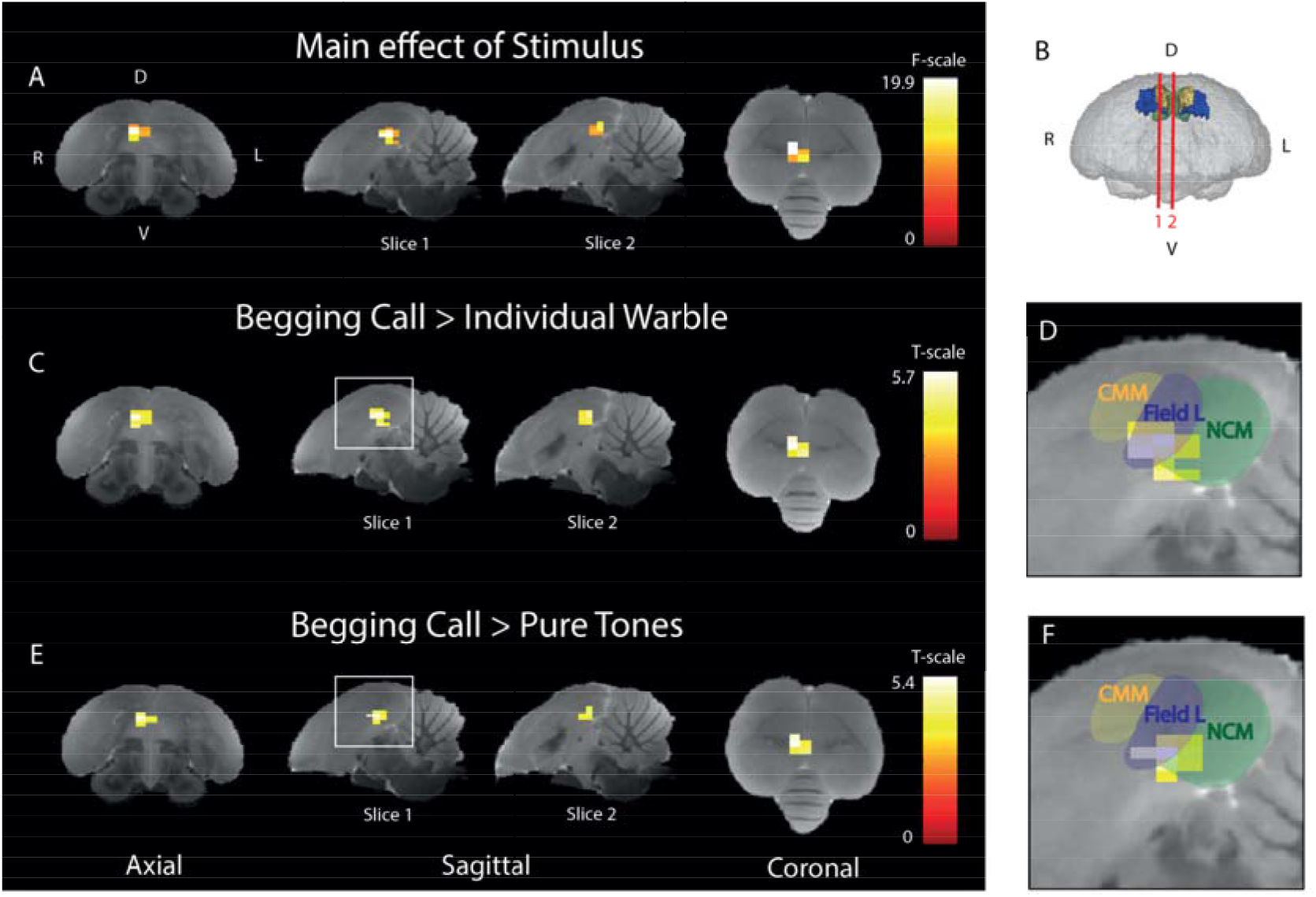
Begging calls sensitivity is highest in right ventral Field L and midsagittal ventral NCM. **(A)** Three orthogonal directions demonstrating the significant main effect of stimulus of the BOLD response. The color bar indicates the F-statistic for each voxel. Only the right (R) hemisphere is depicted as no differential stimulus sensitivity was observed in the left (L) hemisphere **(B)** 3D surface rendering of the starling brain visualizing the slice orientations presented in figures A-F at the level of the auditory forebrain 1.4mm from the midline (slice 1) and 0.16mm from the midline (slice 2). The color bar indicates t-statistics for each voxel. **(C)** Post-hoc test of begging calls (both seasons) compared to individual warbles class 3A (both seasons). The color bar indicates the t-statistics for each voxel. **(D)** Zoom in on peak activation of significant cluster observed in the white square on slice 1 in panel C, overlaid with delineations of CMM, Field L, and NCM. **(E)** Post-hoc test for begging calls (both seasons) compared to pure tones (3 + 7 kHz) (both seasons). The color bar indicates the t-statistics for each voxel. **(F)** Zoom-in on the significant cluster observed in the white square on slice 1 in panel E, overlaid with delineations of CMM, Field L, and NCM. (ANOVA and post-hoc tests; P_FWE_<0.05, k_voxels_ > 10). D=dorsal, V=ventral A=anterior P=posterior, L=left, R=right.

### Begging calls responsiveness is modulated according to seasonal relevance

The next aim was to investigate whether the activation in response to auditory stimulation changes from breeding to non-breeding season. A two-way ANOVA was performed to investigate the main effect of season (**figure 5A**) including both sexes and upon all types of auditory stimuli. A significant seasonal effect was found in ventral NCM covering a cluster of 15 voxels (P_FWE-peak_=0.001, F_peak_=20.86). *Post-hoc* t-tests revealed that the activation is higher in the breeding season in 18 voxels of NCM (P_FWE-peak_=0.001, T_peak_=4.57) compared to the non-breeding season (Figure 5B). No significant voxels were found that displayed a higher activation in the non-breeding season compared to the breeding season.

**Figure 5.**
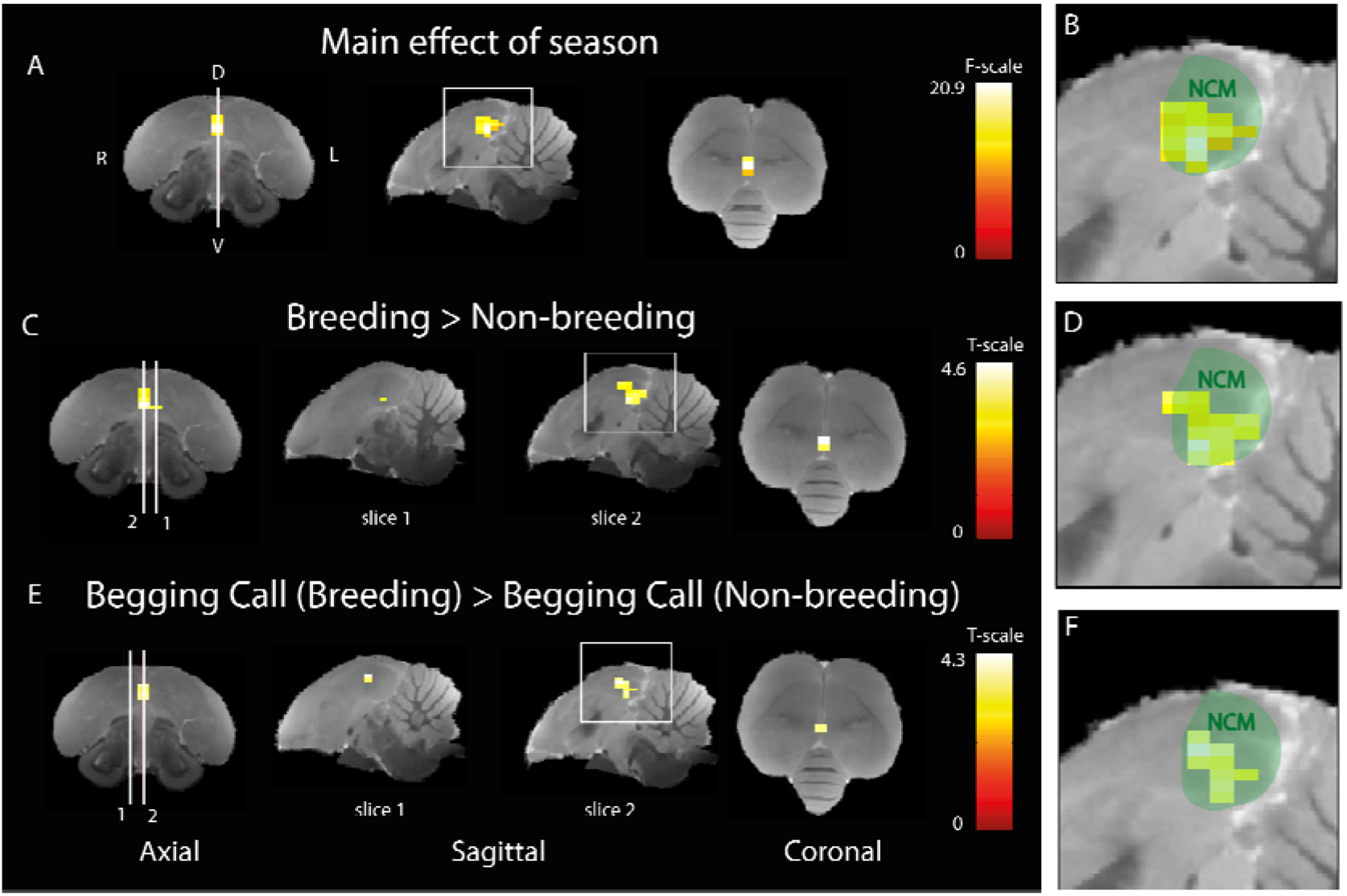
Seasonal difference in the processing of begging calls. **(A)** Three orthogonal slices demonstrating the significant main effect of season (different BOLD responses over rest for breeding compared to non-breeding seasons for all stimuli together. The color bar indicates the F-statistics for each voxel. **(B)** Zoom in on the area in the white square in sagittal view in panel A overlaid on a green delineation indicating NCM. **(C)** Post-hoc comparison of all stimuli during breeding versus non-breeding season. White lines 1 and 2 refer to the position of sagittal slice 1 (1.4mm to the left from the midline) and 2 (0.16mm from the midline) respectively). The color bar indicates the t-statistics for each voxel **(D)** Zoom in on area in the white square in sagittal view in panel C overlaid on a green delineation indicating NCM. **(E)** Post-hoc comparison of differential activation to begging calls stimulation in the breeding versus the non-breeding season. White lines 1 and 2 refer to the position of sagittal slice 1 (1.4mm from the midline) and 2 respectively (0.16mm to the right from the midline). The color bar indicates the t-statistics for each voxel (F) Zoom in on the area in the white square in sagittal view in panel E overlaid on a green delineation indicating NCM. (ANOVA and post-hoc tests; P_FWE_<0.05, k_voxels_ > 10)

Next, we investigated the seasonal differences in activation for the begging calls only by using a t-test including both sexes. A significant increase was found in response to the begging calls in the breeding season compared to the non-breeding season (**figure 5E**; 11 voxels, *P*_FWE-peak_ = 0.002, T_peak_=4.26). The opposite contrast for begging calls in the non-breeding season compared to the breeding season did not result in any significantly different voxels. No significant seasonal differences were found when investigating the individual warbling or the artificial pure tones attesting to our choice of control stimuli and confirming the findings of De Groof et al. (2013 and 2017). Seasonal differences in response to begging calls are located ventrally in the midsagittal region of NCM.

### Sex does not influence the seasonal differential responsiveness to begging calls

Finally, we also investigated if the seasonally different activation for the begging calls in NCM and the intensity of the activation is the same in both sexes. While the two sexes did not respond differently to the begging calls by looking at the main effect of sex, taking together both seasons, the change in perception of the begging calls between seasons may be processed differently by males compared to females. To this end, we tested if there are any voxels where begging calls elicit higher BOLD responses compared to control sounds, in the breeding season vs non-breeding season for males and females separately. This three-way interaction revealed no significant differences in activation patterns between males and females

## Discussion

The current study demonstrates that begging calls produce a significantly stronger and differential activation in comparison with naturalistic and artificial control sounds in right Field L and midsaggital/right NCM. The highest differential activation is found in right Field L showing that already the primary auditory system as such is highly sensitive for this type of acoustic signals that subserve the survival and the fitness of the offspring. Our data also show that the response to begging calls is seasonally modulated in a way that the response is the highest in the breeding season and involves a ventral near-midsagittal part of NCM with the same magnitude in both sexes. These findings corroborate the fact that in starling both parents are actively involved in parental care and at the same time support the existence of an accurate timely functional finetuning of vocal communications between sender and receiver in a breeding context for an important innate vocalization such as the offspring’s begging calls.

### Photoperiodic modulation of hormonal state affects auditory processing of innate calls

The strictly regulated photoperiodic dynamics that were employed in this study to mimic natural seasonality are the main driver for reproduction-related hormonal periodicity in seasonal songbirds (Nicholls, Goldsmith and Dawson, 1988). These light-induced hormonal changes are the main explanation for the demonstrated seasonal differences in how innate vocalizations are perceived and interpreted. Besides the modulation of perception of innate vocalizations, steroid hormones are also known to modulate their production. Exogenous testosterone treatment has been linked to an increase in the *vocal production* of alarm calls in females to match that of males (Gyger *et al*., 1988). and also to enhance nestling begging display, to elicit higher *paternal* feeding behavior (Boncoraglio *et al*., 2006; Noguera, Kim and Velando, 2013). Besides photoperiodic modulation, the hormonal state can be influenced by other factors, such as social structure or attributes, group rearrangements, or the presence of nest boxes. Such factors have a strong effect on sexually dimorphic behaviors and might influence the levels of circulating sex hormones (Riters *et al*., 2000), however, these factors were not present in this study. Other non-hormonal factors such as motivation or attention are also excluded as the birds were mildly anesthetized during the scans. Altogether, the current and previous fMRI studies demonstrate that photoperiod-induced breeding conditions implicate accurate neuromodulations at the level of auditory brain regions to adjust not only sensitivity to relevant learned vocalizations (songs) but also to innate begging vocalizations (calls).

### Processing of learned and innate conspecific vocalizations

ZENK expression studies have revealed that neural activation differs between hearing learned songs versus calls in songbirds and that birds have a preference for conspecific vocalizations over pure tones (Phillmore, Sturdy and Weisman, 2003; Pinaud and Terleph, 2008; Poirier *et al*., 2009). Begging calls in different avian species contain acoustic properties, which reflect the nestling’s condition and species-specific information (Anderson, Brunton and Hauber, 2010; Villain *et al*., 2015; Ursino *et al*., 2018). Processing and categorization of such complex acoustic characteristics are dedicated to higher central auditory regions such as the NCM and CMM. In contrast to field L neurons, CMM and NCM neurons are more selective to a smaller repertoire of natural sounds based on their functional representations (Gentner and Margoliash, 2003). The present study demonstrates that the right NCM seasonally changes its response to innate vocalization. In our previous fMRI work, we have demonstrated that the right NCM can seasonally change its response to *learned* conspecific vocalizations based on their behavioral relevance category in a particular season (De Groof *et al*., 2013, 2017).

We also demonstrated that caudal NCM undergoes microstructural and volumetric changes during the breeding season (De Groof et al., 2009). These seasonal variations in structural and functional properties of caudal NCM coincide with the expression of estrogen receptors and aromatase in songbirds such as the zebra finches (Saldanha *et al*., 2000; Pinaud *et al*., 2006; de Bournonville, Mendoza and Remage-Healey, 2021), white-throated sparrows (Apfelbeck et al., 2013) and canaries (Metzdorf, Gahr and Fusani, 1999), creating a versatile environment for spatiotemporal hormone fluctuations subserving estrogen-dependent modulation of auditory perception (De Groof et al., 2017; see review Spool, Bergan and Remage-Healey, 2022).

Unlike NCM, CMM has weaker associations with estrogenic modulation (Orije and Van der Linden, 2022). Whereas the NCM shows increased activation for novel songs, the CMM is involved in the processing of familiar vocalizations. As the starlings that were used in this study were wild-caught and did not breed in captivity, it is unclear whether they were unfamiliar with begging calls. CMM neurons are also sensitive to social context as they can differentiate between directed and undirected songs (Van Ruijssevelt et al., 2018), and like learned courtship vocalization, begging calls also have directed and undirected renditions(Bulmer, Celis and Gil, 2008; Woolley and Doupe, 2008; Jimeno *et al*., 2014). However, no differences were observed in the CMM between the different stimuli or the seasons.

### Seasonal dynamics restricted to NCM

In a wide variety of aquatic and terrestrial vertebrates, auditory processing is influenced by hormones, even though their precise action mechanisms require further research (*reviewed by* (Hultcrantz, Simonoska and Stenberg, 2006; Canlon and Frisina, 2009; Krentzel and Remage-healey, 2016).

Sisneros and Bass, (2003) demonstrated that seasonal enhancement of auditory perception for the higher harmonic components of the breeding season-specific male-advertisement calls can occur at the peripheral level in the female midshipman fish (*Porichthys notatus*). Even though the peripheral nuclei of the ascending pathway such as the cochlear nucleus (hindbrain), nucleus MLd (midbrain), and nucleus ovoidalis (thalamus) were included in the whole brain field of view of our images, we did not find differences in these regions for any of the stimuli. This gives a strong impression that peripheral processing remains unaffected by seasonality until acoustic signals reach the auditory telencephalon where they cause differential activation.

It is however important to note that these smaller auditory nuclei might be difficult to investigate using auditory fMRI as the signal-to-noise ratio drops noticeably in the ventral parts of the brain, further away from the RF-coil. Besides that, these nuclei (e.g. nucleus ovoidalis and DLM) are often too small to produce detectable BOLD activation that can be included in the average activation mask. While the sensitivity of fMRI is lower in smaller and more ventrally located brain nuclei, it has been demonstrated that fMRI is sensitive enough to pick up subtle differences in song processing at the level of the telencephalon, such as preferences for directed versus undirected song and bird own song over hetero-specific song and even at the level of the midbrain and in a much smaller bird, i.e., the zebra finch (Poirier et al., 2009; van Ruijssevelt et al., 2018).

### The role of sex in auditory perception of begging calls

A study on the spotless Starling (*Sturnus unicolor*), which is closely related to the European starling, revealed a sex bias in their behavioral responsiveness towards begging call playbacks. Only females increase their food provisioning rate in response to nestling begging calls (Kolliker *et al*., 2000; Jimeno *et al*., 2014) even though this species of starlings participate in biparental care (Jimeno *et al*., 2014). One would expect in this case also a sexual difference in the neural substrate responsible for this behavior. However, in our study, no sex differences were found in the way that male and female brains react to begging calls. This suggests that bi-parental care to incubate the eggs as well as in the feeding of the offspring until the fledgling phase of development in avian species, such as the European starling can be the reason why no differences between males and females are observed in the neuronal correlate of hatchlings calls. This is in direct contrast to mammals in which maternal lactation is the primary source of nutrition. It is therefore not surprising that in humans, gender-specific differences have been reported at the level of the amygdala and the orbitofrontal cortex when exposed to infant cry (Seifritz *et al*., 2003; Swain, 2008). In our study, we did not observe activations of the avian amygdala homolog, nucleus taenia (Cheng et al., 1999), but again, the size and location of this nucleus and the use of anesthesia might be a limitation for the fMRI outcome.

### Human sex differences and hormonal modulations in auditory response to infant cries

In humans, the function of the infant cries is similar to that of the avian begging calls. However, in humans, the perception of infant hunger cries has been investigated more extensively with auditory fMRI. From these studies, it becomes clear that activated regions are lateralized towards the right hemisphere and that several brain circuits besides the auditory system are involved. These additional regions include the thalamocingulate circuit (attention shifting), the fronto-insular cortex, the dorsomedial prefrontal cortex and supplementary motor cortex (motor processing and preparation of caregiving response), and lastly the dorsal anterior insula and orbitofrontal cortex (salience detection and evaluation of emotional information) (Witteman *et al*., 2019).

Also, hormonal modulation via exogenous testosterone administration modulates the processing of infant hunger vocalizations in women (Seifritz *et al*., 2003; Bos *et al*., 2010). This suggests that the hormonal modulation of auditory responsiveness to enhance the perception of infant vocalization and its lateralization can be an evolutionarily conserved trait.

## Conclusion

Effective vocal communication requires inducing appropriate neural responses, enabling the execution of relevant behaviors such as mate selection, territorial defense, and parental care. In seasonal songbirds, these particular behaviors are strongly fixed within the boundaries of seasonal reproductive hormone cycles. As a result, hormonal neuromodulation of the auditory system of a receiver is a necessity to match the vocal communication signals of the sender.

We have demonstrated that in starling, highly season-specific innate begging calls elicit more activation in the auditory telencephalon, especially in the NCM, during the breeding season, when begging calls can be expected. This seasonal priming of acoustic perception was shown to be equally important for both sexes supporting the fact that in this species, both parents share an equal role in caring for offspring.

## Acknowledgments

The authors thank Isabel George of the Ethology Animale et Humaine group at the University of Rennes (Rennes, France) for providing them with starling warbles and begging recordings that were used in this research. This research was supported by a postdoctoral fellowship awarded to EJ (grant agreement no. 12R1917N) and a research project awarded to AVdL (grant agreement no. G030213N) both from the Research Foundation Flanders (FWO). The computational resources and services used in this work were provided by the HPC core facility CalcUA of the Universiteit Antwerpen, and VSC (Flemish Supercomputer Center), funded by the Research Foundation - Flanders (FWO) and the Flemish Government. The equipment used in this study was supported by the Hercules stichting (grant agreement no. AUHA/012) awarded to AVdL, financing MRI-research infrastructure.

## Appendix

**Supplementary figure 1.**
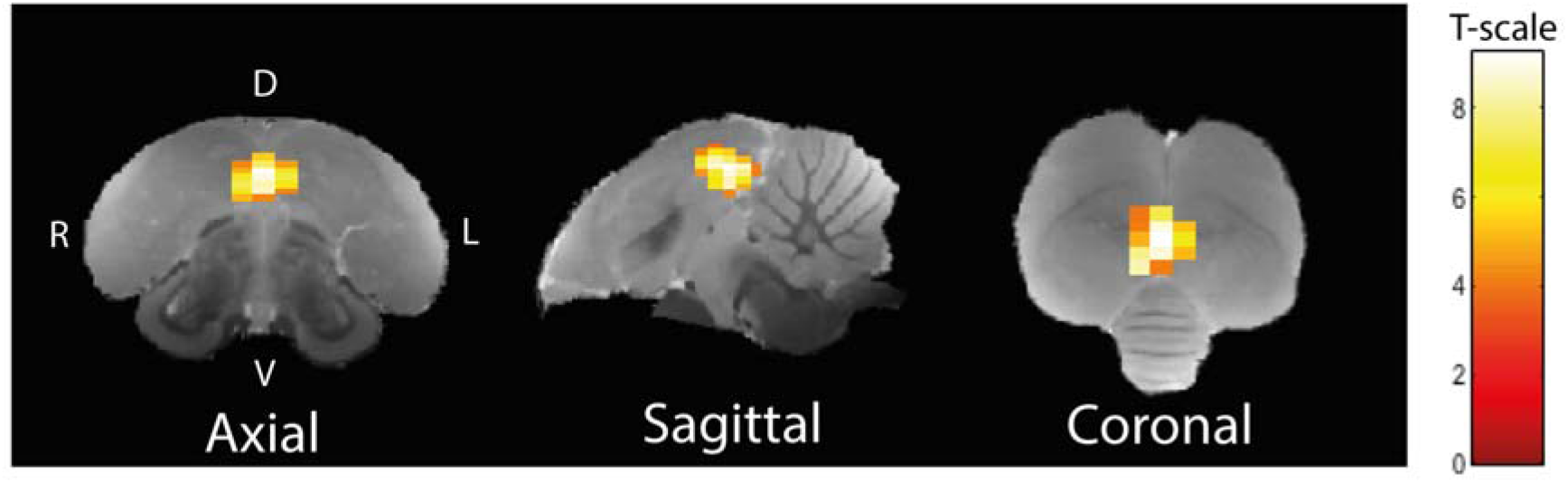
One sample T-test demonstrating significant (P_UNC_<0.001, k_voxels_ > 10) activation of average auditory stimulation block (begging call, individual warble, and pure tones) over rest periods. Color bars indicate a significant BOLD-response higher than rest periods.

